# APOC2 accelerates colorectal cancer progression via modulating lipid metabolism and transcriptional activity of PHF8

**DOI:** 10.1101/680967

**Authors:** Jian Chen, Chao Xiao, Yupeng Wang, Guohe Song, Xiaoliang Wang, Xueni Liu, Jiayi Chen, Huijun Lu, Jing Kuai, Xuebin Qin, Weiping Guo, Huamei Tang, Zhihai Peng

## Abstract

The apolipoproteins (APOs) are the major proteins in blood lipid transportation. Emerging evidence has demonstrated that APOs might exert important function in tumor cells, but the underlying mechanism remains inclusive. In this study, we aim to explore the relationship between APOC2 dysfunction and colorectal cancer (CRC) malignancy. By analyzing the expression of APOC2 in 507 patients with CRC, we demonstrated that the APOC2 was overexpressed and associated with poor prognosis in CRC. We then found that high levels of APOC2 resulted in proliferation, invasion and metastasis of CRC cells in vitro and in vivo. Mechanistically, we revealed that APOC2 directly interacted with FASN which resulted in decreased levels of omega-3 fatty acids and increased levels of alpha-ketoglutarate (α-KG). Both RNA-seq and ChIP-seq analysis revealed that APOC2 overexpression resulted accumulation of α-KG leads to activation on the transcriptional program of PHF8 and thereby contributed to activation on genes involved in cell proliferation, invasion and metastasis. Together, our study unveiled the oncogenic role of APOC2 in tumor cells, which sheds new light on the potential of APOC2 as a biomarker in the prognosis of CRC.

## Introduction

Colorectal cancer (CRC) is one of the most lethal malignancies in the world[1]. Due to the lack of early specific symptoms[2], early-stage CRC is difficult to identify. Emerging evidence indicates that a lipid metabolic disturbance is a critical risk factor in the occurrence and development of colorectal cancer[3, 4]. Reprogramming of lipid metabolism in cancer cells is crucial for providing energy and substrates for biosynthesis, which are required for cancer cell proliferation and invasion[5, 6]. In addition, metabolites generated from lipid metabolism, including alpha-ketoglutarate, acetate and leukotrienes, may themselves be involved in cancer development[7, 8].

Apolipoproteins are the major lipoproteins involved in lipid transport, and are mainly expressed in liver tissue and delivered through blood vessels[9, 10]. Dysregulation of apolipoproteins is involved in many diseases including vascular disease, heart disease, Alzheimer disease, and cancers. Recent studies show that dysregulation of serum apolipoproteins highly correlates with distinct outcomes of cancer therapy[10–15]. A high level of apolipoprotein A1 is associated with cancer risk while a low level of apolipoprotein B is associated with risk of breast cancer[10]. Apolipoproteins in the blood transfer lipids to cancer cells to provide energy for cancer cell proliferation and invasion. Apolipoproteins also function as important factors in cellular signal transduction. For instance, apolipoprotein E (APOE) interacts with low density lipoprotein receptor-related protein 1 (LRP1) and low-density lipoprotein receptor-related protein 8 (LRP8) in tumor cells and inhibits tumor cell invasion[16]. Apolipoprotein B (APOB) represses angiogenesis, a key process for tumor cell invasion and metastasis, by modulating expression of the VEGF receptor[17]. Apolipoprotein C1 is involved in maintaining cell survival of pancreatic cancer cells through the regulation of cell apoptosis[18].

Apolipoprotein C2 (APOC2), an important member of the apolipoprotein gene family, is primarily biosynthesized in the liver and intestine, and is present on plasma chylomicrons, VLDL, and high-density lipoproteins (HDL). It activates the enzyme lipoprotein lipase, which hydrolyzes triglycerides and thus provides free fatty acids for cells[19, 20]. Defects in the structure or production of APOC2 and some other gene products result in conditions of LPL deficiency such as hypertriglyceridemia and chylomicronemia [21]. Recent studies show that high serum APOC2 levels are correlated with poor prognosis of pancreatic adenocarcinoma. High levels of APOC2 promote pancreatic cancer cell growth and invasion[22], suggesting a potential oncogenic role for APOC2 during tumorigenesis, but the underlying mechanism remains unclear. Of note, the majority of studies on apolipoprotein are focused on its levels in blood, the roles of apolipoproteins in tumor cells remain largely unclear.

Here in our study, we observed that APOC2 is overexpressed in colorectal cancer cells and associated with poor prognosis. We found that overexpression of APOC2 accelerates tumor progression and metastasis of CRC. APOC2 directly interact with FASN and leads to reduced omega-3 fatty acids in CRC cells and results in accumulation of alpha-ketoglutarate acid (α-KG). The accumulated α-KG activates the transcriptional programs of PHF8. Furthermore, we demonstrated the activation of APOC2 is caused by DNA hypo-methylation and transcriptional activation by MYC and E2F1.

## Results

### APOC2 is overexpressed and highly correlates with poor prognosis in colorectal cancer

To investigate the expression of APOC2 in colorectal cancer, we used real-time quantitative PCR (RT-qPCR) to compare APOC2 transcript levels in colorectal cancer (CRC) tissue versus corresponding adjacent normal tissue in 32-paired patient samples. We found that 20 of 32 patients showed over 2-fold higher APOC2 expression in CRC tissue compared to adjacent normal tissue. (Fig. 1A). We also compared APOC2 expression levels in CRC tissues and adjacent normal tissues in three different datasets from the Genome Expression Omnibus (GEO). We found that CRCs had significantly higher levels of APOC2 than normal colon tissues (Fig. 1B; **Fig. S1**). We next used immunohistochemical staining to evaluate APOC2 expression in patients obtained from two independent clinical cohorts (Fig. 1C). The cohorts 1 and 2 comprised paired adjacent normal colon and CRC samples of 276 patients collected from 2003 to 2007 and of 231 patients collected from 2008 to 2011, respectively. We demonstrated that APOC2 expression in CRC was significantly higher than that in normal tissue in paired samples from both cohort 1 and cohort 2 (Fig. 1D). Together, these findings indicate that APOC2 is highly expressed in CRC. To evaluate the potential prognostic value of APOC2 expression, we investigated whether APOC2 expression is correlated with the prognosis and/or clinical characteristics of CRC patients. We performed multivariate analysis in cohort 1 and 2 and found that high expression of APOC2 was significantly associated with poor survival in patients with CRC independently of the AJCC stage (Fig. 1E and G; **Table S1**). Kaplan-Meier analysis in both cohorts showed that a high level of APOC2 expression was associated with poor disease-free survival (DFS) and overall survival (OS) in both cohorts (Fig. 1F and H; **Fig. S2**). We further analyzed the correlation between APOC2 expression and clinical outcomes using the dataset from The Cancer Genome Atlas (TCGA). Kaplan-Meier survival curves showed significantly shorter DFS in CRC patients with high APOC2 gene expression compared to patients with low expression (Fig. 1I). These data indicate that APOC2 expression could serve as a novel prognostic factor in patients with CRC, and also suggest a potential oncogenic function of APOC2 in CRC.

**Fig. 1.**
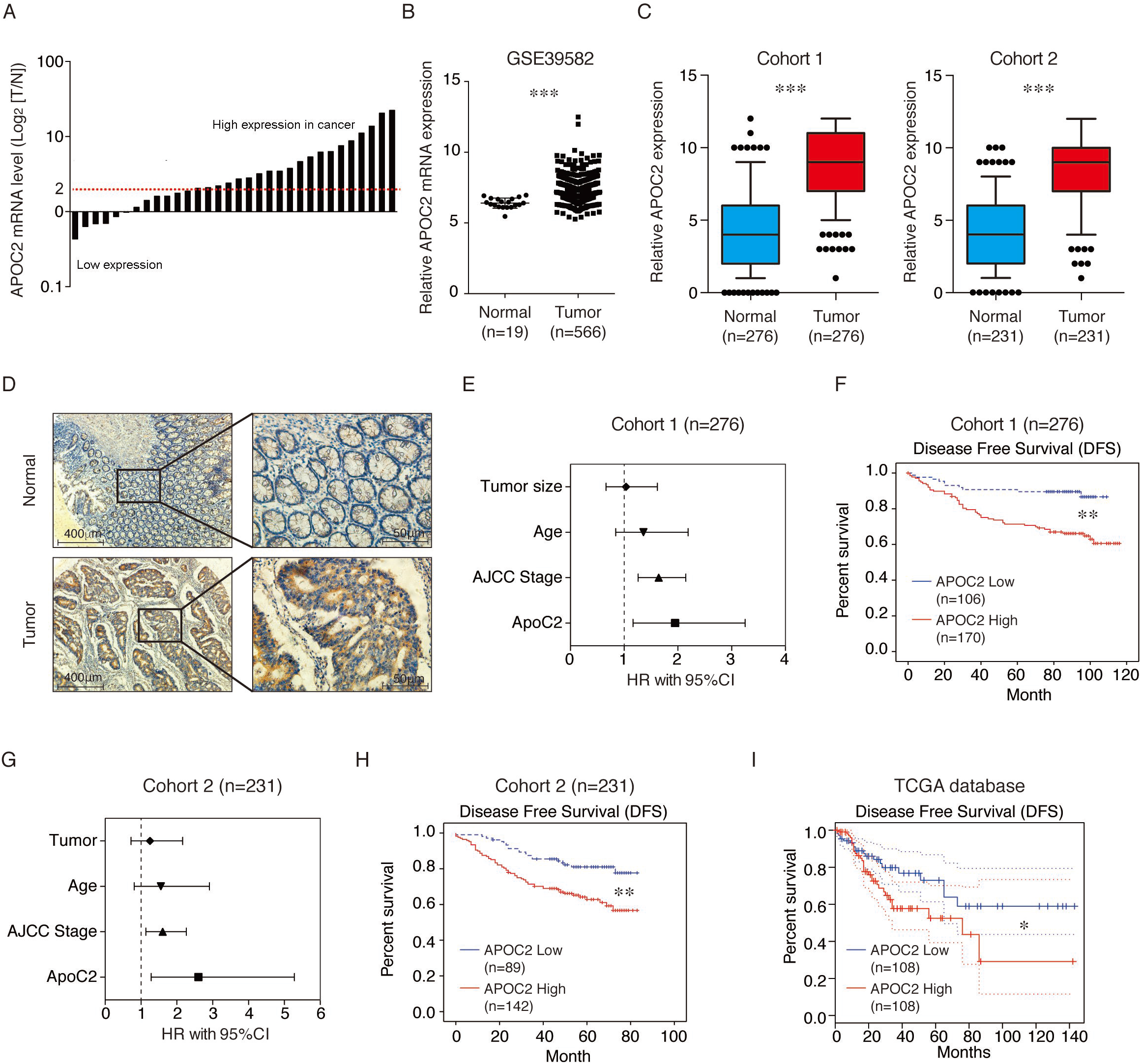
APOC2 is overexpressed and highly correlates with poor prognosis in colorectal cancer. (**A**) APOC2 expression in colorectal cancers and their matched adjacent normal tissues. Waterfall plot show that APOC2 mRNA expression is highly expressed in tumors as compared to adjacent normal colon tissues. (**B**) Analysis of APOC2 expression in CRC and their adjacent normal tissues using dataset GSE39582 from GEO database. (**C**) Box plots illustrate the relative expression of APOC2 in matched normal colorectal tissue and CRC in two independent cohorts. The horizontal lines represent the medians; the boxes represent the interquartile range, and the whiskers represent the 10^th^ and 90^th^ percentiles. *** *p* < 0.001, Student’s t-test. (**D**) Representative immunohistochemical staining for APOC2 in CRC (bottom panel) and adjacent normal colorectal tissue (upper panel). APOC2 expression in 507 paired samples of CRC and normal colon tissue was evaluated by IHC. (**E** and **G**). Survival analysis showed that high APOC2 expression is associated with poor prognosis in CRC patients from cohort I (E) and cohort II (G). Log-rank test. (**F** and **H**). Disease-Free Survival (DFS) was compared between patients with low and high expression of APOC2 in cohort I (F) and cohort II (H). (**I**) DFS was compared between CRC patients with low and high APOC2 expression from the TCGA database.

### APOC2 is correlated with recurrence and metastasis of colorectal cancer

Since recurrence is the major cause of poor prognosis, we next sought to analyze the expression of APOC2 in primary tumors of patients with recurrence or without recurrence in our cohorts. Recurrence was found in 85 patients of cohort 1. In cohort 2, 63 patients had CRC recurrence after surgery (**Table S1)**. Patients with recurrence in our cohorts, as well as such patients from the TCGA database, had significantly higher levels of APOC2 expression than patients without recurrence (Fig. 2A-C). We further analyzed the expression level of APOC2 in primary tumor cells and metastasized tumor cells in lymph nodes from our two cohorts and from the GEO database. APOC2 expression in the metastasized tumor cells was much higher than that in paired non-metastasized tumor cells (Fig. 2D-F), These data suggest that high APOC2 expression may also be associated with CRC metastasis.

**Fig. 2.**
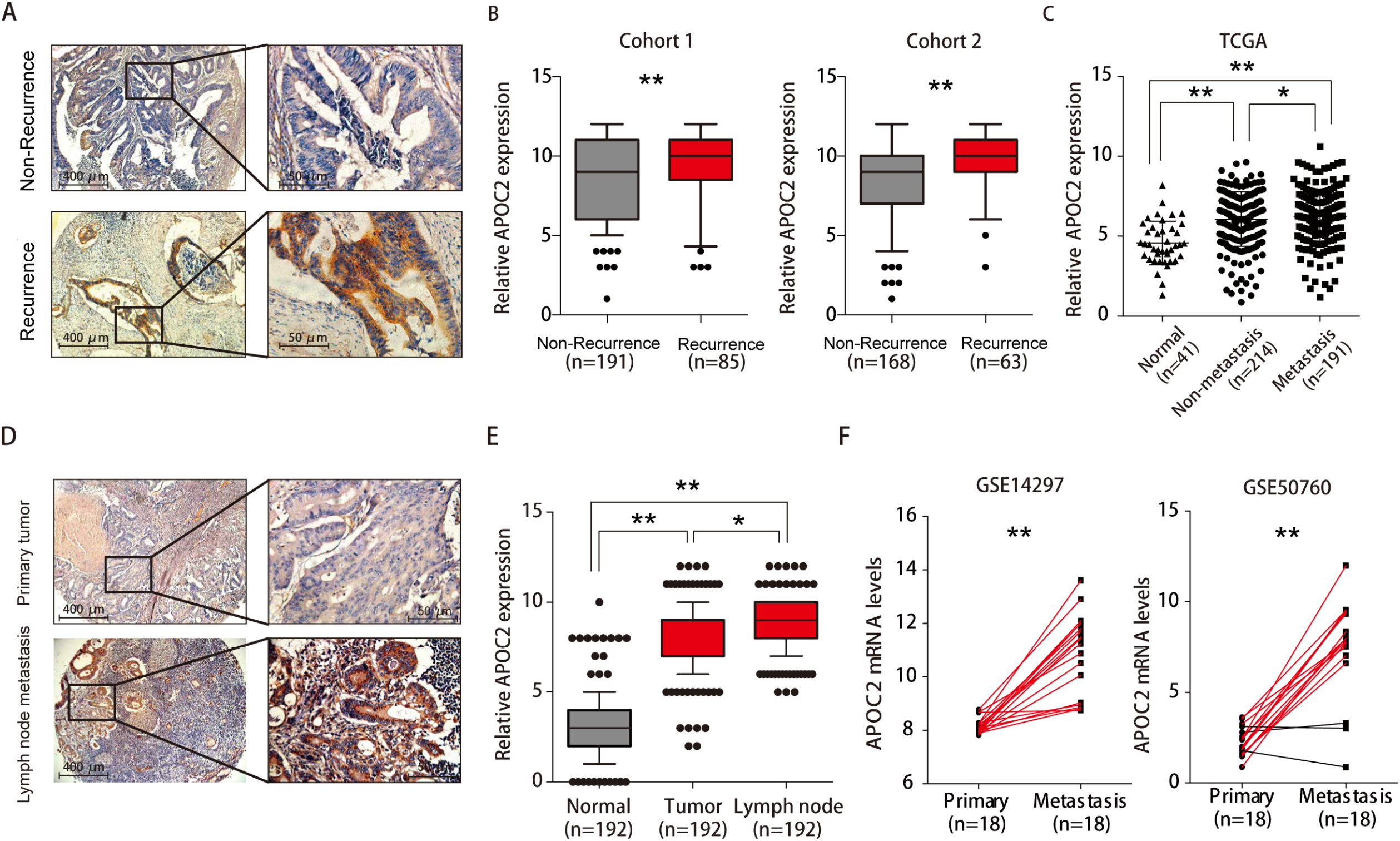
APOC2 is correlated with recurrence and metastasis of colorectal cancer. (**A**) Representative immunohistochemical staining of APOC2 proteins in CRC tissues from patients without recurrence (upper) and with recurrence (lower). (**B**) Statistical analysis of the expression of APOC2 proteins in CRC tissues from patients without recurrence (upper) and with recurrence (lower) from two independent cohorts. (**C**) Statistical analysis of the expression of APOC2 proteins in CRC tissues from patients without metastasis and with metastasis or in normal tissues from the TCGA database (**D**) Representative immunohistochemical staining of APOC2 proteins in primary CRC tissues (upper) and in metastasized tumor cells in lymph nodes (lower). (**E**) Statistical analysis of the expression of APOC2 proteins in normal tissues, CRC tissues, and metastasized tumor cells in lymph nodes. (**F**) Analysis of APOC2 expression in primary CRC tissues and in metastasized tumor cells using datasets from GEO databases.

### APOC2 promotes colorectal cancer cell proliferation and invasion *in vitro*

To investigate the role of APOC2 in colon cancer tumorigenesis and metastasis, we altered expression levels of APOC2 in cultured CRC cells and tested the effect on clone formation ability, cell proliferation, migration, and invasion. APOC2 was overexpressed in Caco2 and RKO cells lines, which normally express low levels of APOC2. APOC2 expression was knocked down in SW620 and HT29 cell lines, which normally express high levels of APOC2 (**Fig. S3**). We found that overexpression or inhibition of APOC2 expression resulted in increased or decreased clone formation (Fig. 3A and B), cell proliferation (Fig. 3C and D), migration (Fig. 3E and F) and invasion (Fig. 3G and H), respectively. Together, these findings regarding the promotion of cell proliferation and migration mediated by APOC2 suggest that APOC2 may function as an oncogene in colorectal cancer.

**Fig. 3.**
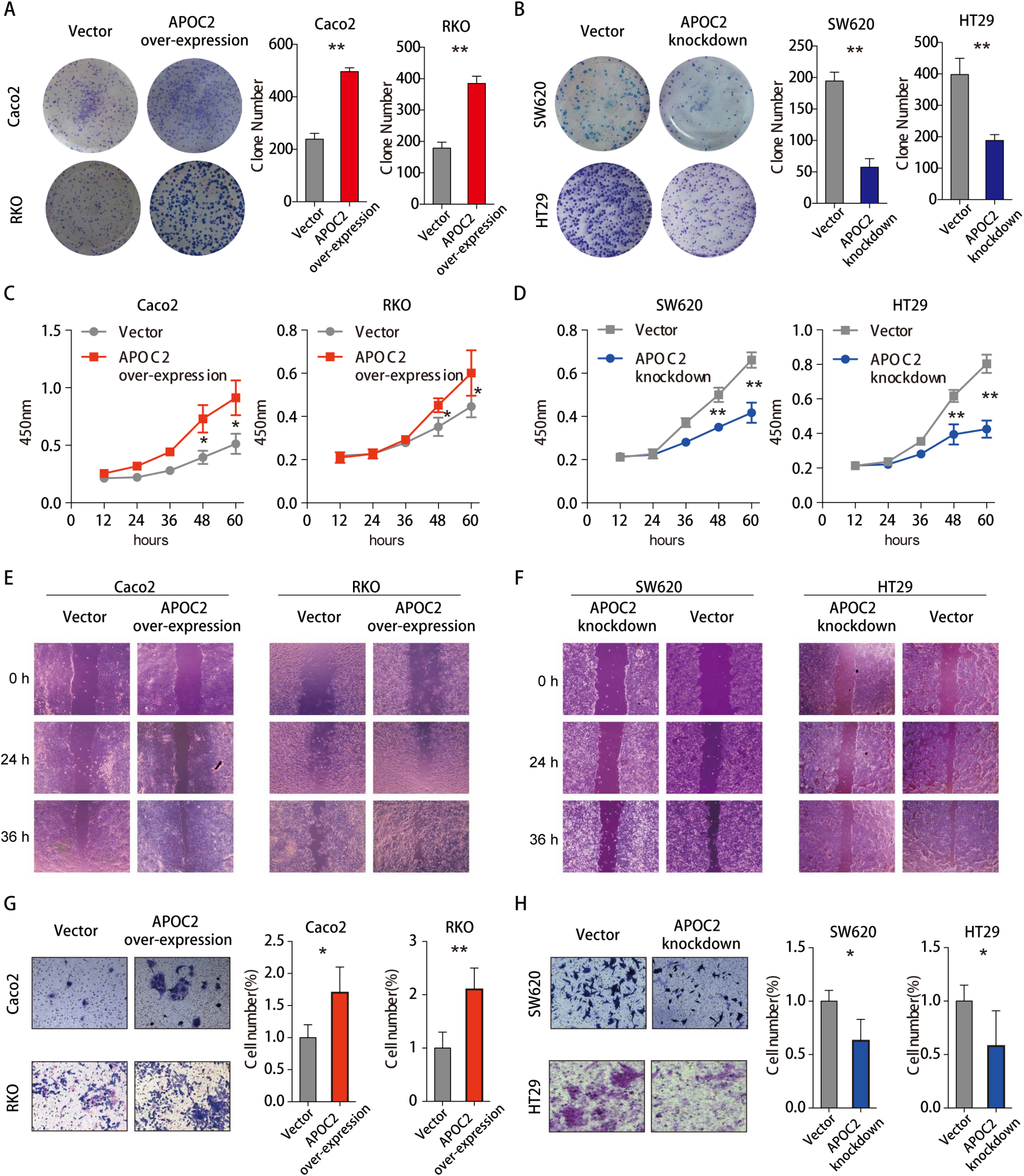
APOC2 promotes colorectal cancer cell proliferation and invasion *in vitro*. (**A** and **B**) Colony formation assays using cells overexpressing APOC2, APOC2 knockdown cells, or cells transfected with vector controls. Caco2 and RKO cells were transfected with control vector or APOC2 overexpression plasmids. SW620 and HT29 cells were transfected with control vector or APOC2 shRNA plasmids. Images of the whole plate are shown. (**C** and **D**) Cell proliferation was assessed by CCK8 assay. Cell viability was evaluated at an optical density of 450 nm. Caco2 and RKO cells were transfected with the control vector or APOC2 overexpression plasmids. SW620 and HT29 cells were transfected with control vector or APOC2 shRNA plasmids. Data are means ± SD. (**E-G**) Cell motility was analyzed by wound-healing assays and invasion assays. Caco2, RKO, SW620 and HT29 cells were stably transfected with the indicated plasmid. Data shown are representative of three independent experiments. Data are means ± SD. \**p* < 0.01.

### APOC2 promotes colorectal cancer progression and metastasis *in vivo*

To assess the effects on CRC of APOC2 *in vivo*, we generated APOC2 stably overexpressing and knockdown cells and injected them into the flanks of nude mice. As illustrated in Figure 4A-D, APOC2 overexpression increased tumor size and volumes while knockdown of APOC2 resulted in decreased tumor size and volumes. This indicates that APOC2 substantially promotes cancer progression of CRC *in vivo*.

**Fig. 4.**
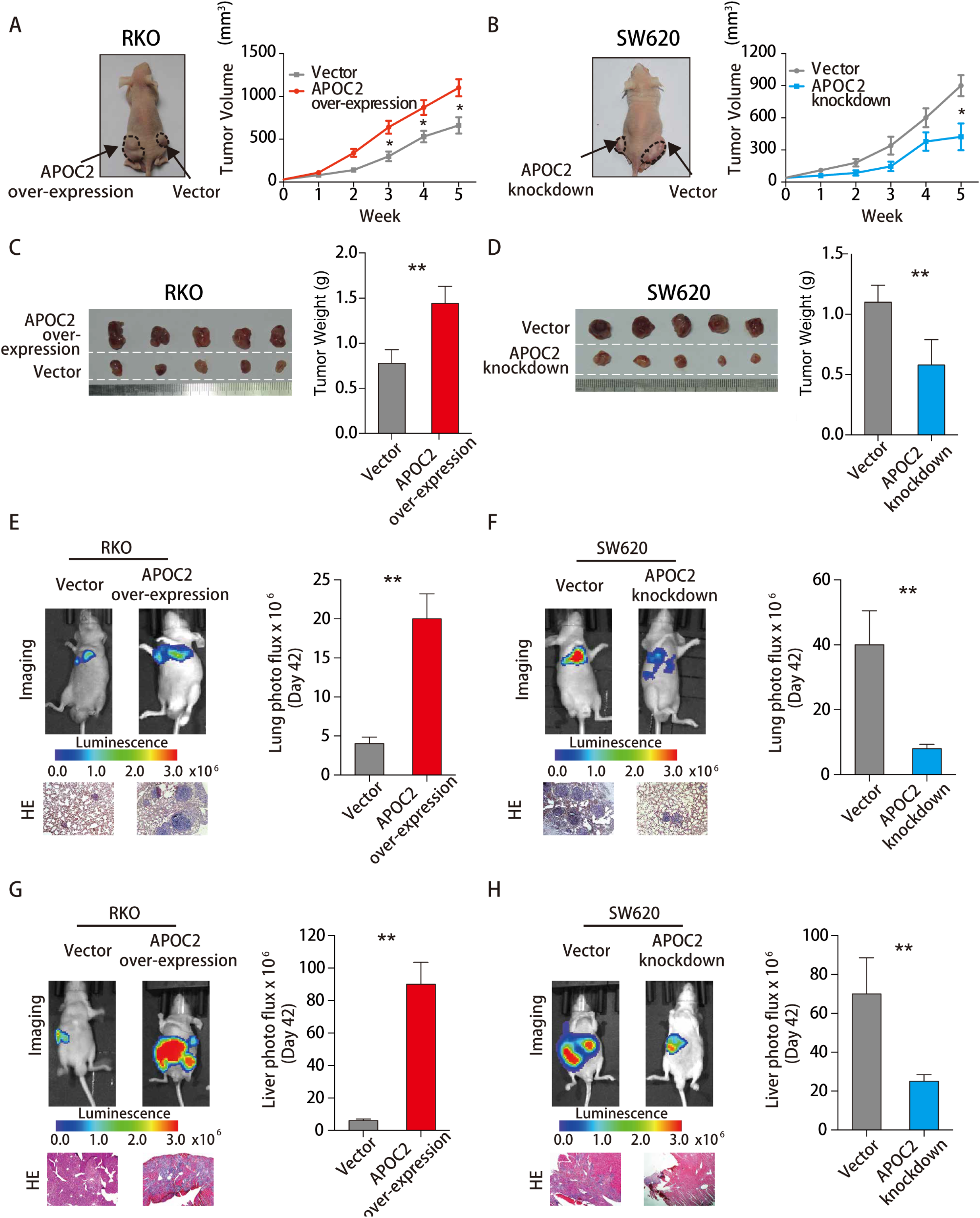
APOC2 promotes colorectal cancer progression and metastasis *in vivo*. (**A**) APOC2 accelerated tumor growth of RKO cells in nude mice. APOC2 overexpressing RKO cells or vector infected RKO cells were injected into nude mice (n = 6). Tumor growth curves with time series are displayed. (**B**) Knockdown of APOC2 inhibited tumor growth of SW620 cells in nude mice. Tumor growth curves with time series are displayed. Data are means ± SD. (**C**) Increased tumor size with APOC2 overexpression. (**D**) Decreased tumor size with APOC2 knockdown. The tumors were isolated at the end of the assay, and the tumor weights of each group were measured. Data are means ± SD. (**E**) Overexpression of APOC2 increased CRC lung metastasis. (**F**) Knockdown of APOC2 decreased CRC lung metastasis. (**G**) Overexpression of APOC2 increased CRC liver metastasis. (**H**) Knockdown of APOC2 decreased CRC liver metastasis. (E-H) Luciferase stably expressing RKO cells were transfected with APOC2 overexpression plasmid or vector control, and luciferase stably expressing SW620 cells were transfected with APOC2 knockdown plasmid or vector control. Representative bioluminescence images of mice injected with indicated CRC cells are shown in the left panel of each subfigure.

To directly monitor the effect of APOC2 on CRC cell growth and metastasis *in vivo*, we co-expressed a luciferase reporter plasmid in the APOC2 overexpressing or APOC2 knockdown cell lines to label each cell individually. These Luc-labelled lines were xenografted into nude mice. Tumor metastasis were respectively studied by subcutaneous injection into nude mice by either tail vein (for lung metastasis) or local spleen injections (for liver metastasis). The growth of tumors was measured every week, and xenograft tumors were removed for final analysis. CRC lung metastasis and liver metastasis were detected by fluorescent photon fluxes, respectively. We found that overexpression or knockdown of APOC2 resulted in increased or decreased metastasis (Fig. 4E-H), respectively. Together, our *in vivo* data demonstrate that APOC2 promotes tumor cell progression and metastasis in colorectal cancer.

### APOC2 regulates cellular lipid metabolism and interacts with FASN in CRC

APOC2 is an apolipoprotein which normally functions in the transport of lipids to tissues and which otherwise participates in lipid metabolism. Our finding that overexpression of APOC2 is associated with tumorigenesis and metastasis prompted us to identify the cellular lipid metabolic products associated with APOC2 in CRC. We performed quantitative analysis of free fatty acids via LC/MS (liquid chromatography–mass spectrometry) analysis in APOC2 overexpressing RKO cells (**Table S2**). We found that the fatty acids which showed the most significant expression changes between vector-control and APOC2 transfected cells were eicosapentaenoic acid (EPA) and docosahexaenoic acid (DHA) (Fig. 5A). As these two lipid products have an anti-cancer effect on many tissues[23–25], it is conceivable that decreased DHA and EPA resulting from overexpression of APOC2 may contribute to tumorigenesis. Indeed, addition of either EPA or DHA to the high APOC2-expressing cell lines SW620 and HT29 suppressed cell proliferation (Fig. 5B and C). This result indicates that inhibition of DHA and EPA levels by APOC2 benefits CRC cell growth. Thus, the oncogenic effects of APOC2 may be mediated at least partly through regulation of DHA and EPA levels.

**Fig. 5.**
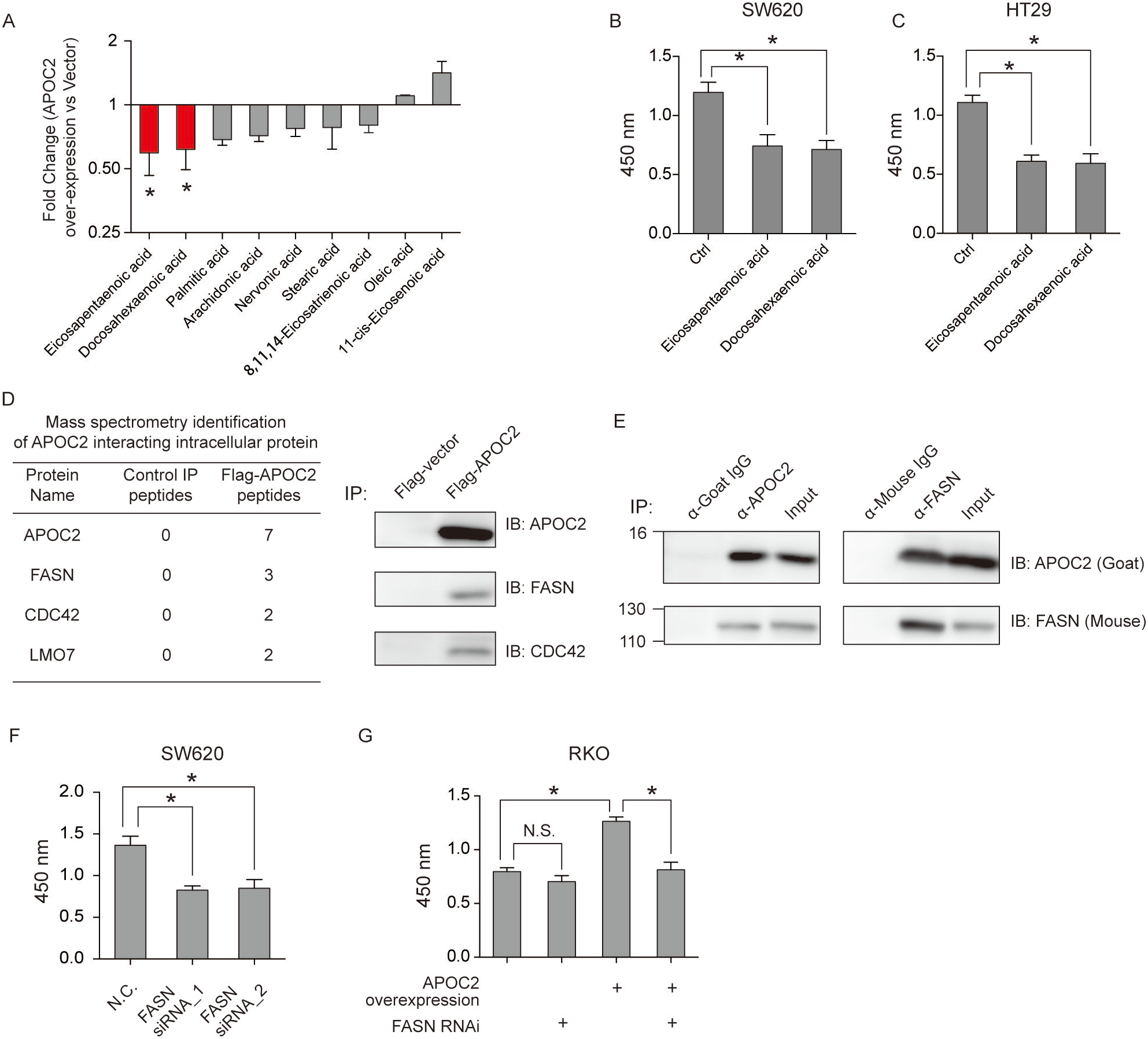
APOC2 regulates cellular lipid metabolism and interacts with FASN in CRC. (**A**) Changes in fatty acid levels in RKO cells with or without APOC2 overexpression. Cellular fatty acid levels were evaluated by mass spectrometry analysis. (**B** and **C**) Effect of eicosapentaenoic acid (EPA) and docosahexaenoic acid (DHA) on SW620 cells (B) and HT29 (C) cells. Both cells were treated with EPA or DHA for 48 hours. Cell viability was evaluated by CCK8. Data are means ± SD. (**D**) APOC2 is associated with several lipids metabolism--associated proteins, as shown by mass spectrometry analysis after pulldown of Flag-tagged APOC2. Validation of APOC2 interaction with proteins was validated by immunoprecipitation followed by Western blot. RKO cells were transfected with Flag-tagged APOC2 and immunoprecipitation were performed using Flag gel beads. The eluted protein complex was subjected to Western blot using indicated antibodies. (**E**) Interaction of APOC2 with FASN in SW620 cells. Co-immunoprecipitation experiments were performed in SW620 cells using the FASN antibody and APOC2 antibody. (**F**) Knockdown of FASN decreased growth of SW620 cells. Two siRNAs targeting FASN were transfected into SW620 cells for 48 hrs, and cell viability was evaluated by CCK8 assays. (**G**) APOC2-mediated cell growth was blocked by knockdown of FASN expression. Cells stably transfected with APOC2 or vector were treated with FASN siRNA for 48 hrs. Cell viability was evaluated by CCK8 assays.

Since APOC2 does not have enzymatic activity to degrade lipid products such as EPA and DHA. To understand how APOC2 is able to regulate these lipid levels, we conducted proteomic analysis of cells overexpressing APOC2 and corresponding control cells. We overexpressed Flag-tagged APOC2 in RKO cells (which normally express low levels of APOC2) and performed immunoprecipitation followed by LC-MS/MS to identify the cellular partners of APOC2 in CRC. As shown in Figure 5D, APOC2 interacts with FASN (fatty acid synthase) in CRC. FASN is one of the key regulators of cellular lipid metabolism and links lipid metabolism to cell growth and invasion. Co-immunoprecipitation using antibodies against APOC2 and FASN in SW620 cells (Fig. 5E) showed that FASN was specifically detected in the protein complex immunoprecipitated by APOC2-specific antibodies but not in the protein complexes precipitated by the normal IgG control in SW620 cells. Similarly, APOC2 was specifically detected in the protein complexes immunoprecipitated by FASN-specific antibody. Thus, APOC2 may regulate DHA and EPA levels via FASN.

To determine whether APOC2 contributes to CRC progression through the APOC2-FASN axis, we overexpressed APOC2 in CRC cells with or without knockdown of FASN expression. As shown in Figure 5F and 5G, suppression of FASN expression significantly reduced the acceleration of cell growth mediated by APOC2.

### APOC2 regulates the expression of oncogenes involved in cellular metabolism, cell cycle and MAPK cascade

Recent studies indicate that apolipoprotein E (APOE) and apolipoprotein B (APOB) contribute to tumorigenesis through directly binding with critical downstream proteins such as LRP1/LRP8 and VEGF receptor[16, 17], respectively. These findings prompted us to further explore the downstream events by which APOC2 participates in tumorigenesis and tumor metastasis in *vitro*. By microarray gene analysis of vector-control and APOC2 overexpressing RKO cells, we identified 4,322 genes differentially expressed upon APOC2 overexpression (**Table S3**). We found that the majority of these genes were upregulated, suggesting that APOC2 may be involved in transcriptionally upregulating genes in RKO cells. We also performed gene ontology enrichment analysis on this identified set of genes. We found that many of these genes are involved in processes related to cell growth and metastasis in CRC, including cellular metabolism, the MAPK pathway, and the cell cycle (Fig. 6A and B). This result is consistent with observations obtained from gene set enrichment analysis (GSEA) using the RNA-seq datasets from the TCGA database (Fig. 6C).

**Fig. 6.**
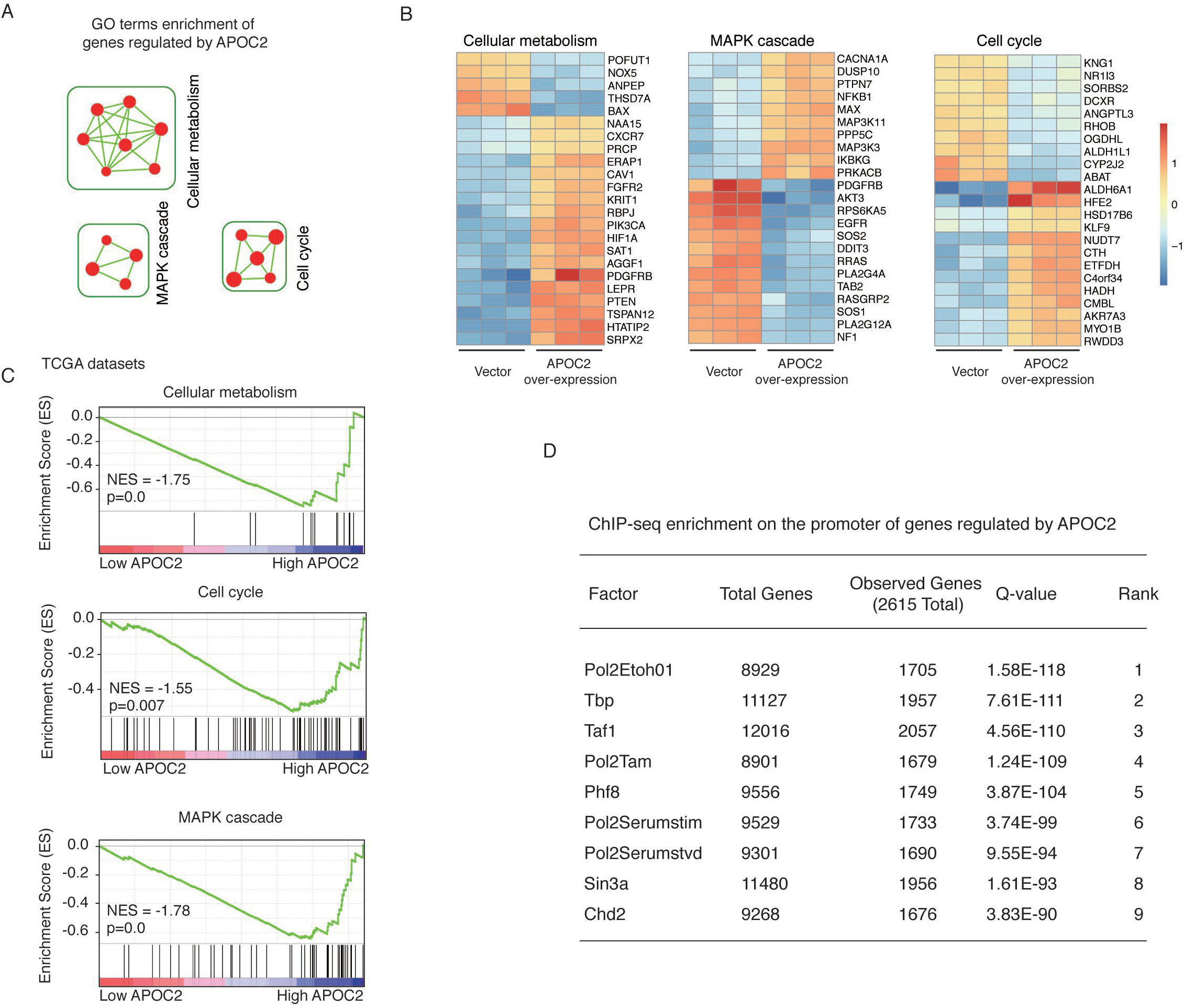
APOC2 regulates the expression of oncogenes involved in cellular metabolism, cell cycle and MAPK cascade. (**A**) Gene ontology enrichment analysis of genes regulated by APOC2. Gene ontology enrichment map showing that genes regulated by APOC2 are involved in the cell cycle and MAPK cascade as well as in cellular metabolism. Nodes represent individual GO terms and edges represent the degree of relatedness between connecting GO terms. The indicated clusters were identified using the Enrichment Map. (**B**) Heatmaps showing the representative GO terms enriched within APOC2 regulated genes. (**C**) GSEA analysis showing the correlation between the gene signatures specific to APOC2 overexpression and genes involved in the cell cycle, MAPK cascade, and cellular metabolism. The gene expression profiles of CRC patients were retrieved from TCGA. NES, normalized enrichment score. The expression level of representative genes was calculated and plotted as log2(RPKM+1). (**D**) Genes upregulated by APOC2 were enriched in binding sites for PHF8. The ENCODE ChIP-seq significance tool were used to analyze the enrichment of specific transcription factor binding sites in the promotors of genes upregulated by APOC2

To further explore the mechanism by which APOC2 regulates the transcription of these genes in CRC, we performed transcription factor enrichment analysis using the ChIP-seq data from the ENCODE project and the genes differentially expressed upon APOC2 overexpression. We found that the genes activated by APOC2 were significantly enriched in genes bound by PHF8 (Fig. 6D). This suggests that PHF8 may directly regulate these APOC2-activated genes.

### APOC2 activates downstream oncogenes by modulating the transcriptional activity of PHF8

To evaluate the prediction that PHF8 may directly regulate these APOC2-activated genes, we performed ChIP-seq using antibodies against PHF8 in APOC2 overexpressing and control RKO cells. We identified 4,015 targets in control cells and 8,928 targets in APOC2 overexpressing RKO cells (Fig. 7A and **Table S4**) and found that the mean PHF8 ChIP-seq signal of APOC2-overexpressing cells was evident compared with that of control (Fig. 7B). We identified MAPK1, DAD1, NFKB1, CDK2, MYO1B and ACACA, all proteins with established roles in tumorigenesis, as proteins which had stronger binding activity to PHF8 in APOC2 overexpressing cells than in control cells (Fig. 7C). These results indicate that APOC2 indeed affects binding of PHF8 on chromatin in CRC.

**Fig. 7.**
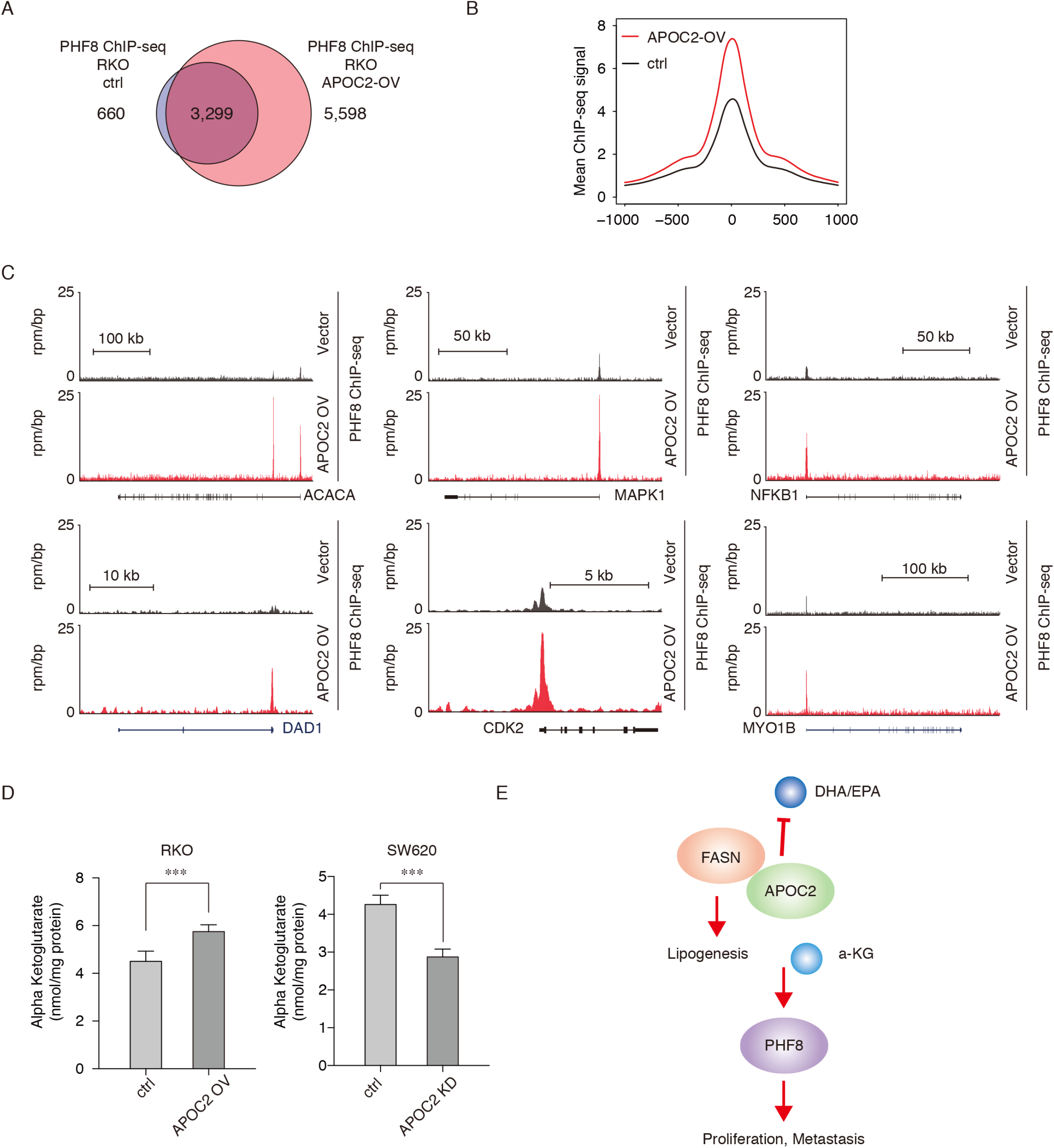
APOC2 activates downstream oncogenes by modulating the transcriptional activity of PHF8. (**A**) APOC2 induced increased targets of PHF8 on chromatin. Venn diagram of the overlap of PHF8 binding sites with or without APOC2 overexpression in RKO cells. were illustrated. (**B**) Metagene representations of the average ChIP-seq signal for PHF8 across the PHF8 binding sites with or without APOC2 overexpression. Each metagene is centered on the PHF8 summit. (**C**) Increased binding of PHF8 at the promoter of APC. ChIP-seq binding profiles for PHF8 at the regulatory regions of representative genes are presented. ChIP-seq was performed in RKO control cells and in APOC2 overexpressing RKO cells. (**D**) Correlation between APOC2 expression and cellular α-KG levels. (**E**) Model showing the role of APOC2 in colorectal cancer. APOC2 overexpression leads to increased DHA and EPA usage, thereby affecting the level of cellular α-KG. α-KG directly affects PHF8 binding on chromatin and leads to activation of genes involved in cell proliferation, cell invasion, and the MAPK cascade.

Alpha-ketoglutaric acid (α-KG) is a known key co-factor for PHF8 and is essential for PHF8 to regulate expression of downstream target genes [26–28]. To investigate the mechanism underlying PHF8 activation associated with APOC2, we measured the level of α-KG in APOC2 overexpressing RKO cells and in APOC2 knockdown SW620 cells, as well as in the respective control cells. As shown in Figure 7D, α-KG levels were significantly increased with APOC2 overexpression and reduced with APOC2 knockdown. These results indicate that APOC2 expression is related to the production of α-KG in cancer cells. The level of α-KG may affect the transcriptional activity of PHF8, thereby leading to transcriptional upregulation of genes involved in CRC proliferation and metastasis (Fig. 7E).

### DNA hypo-methylation and the transcriptional activation by MYC and ELF1 contribute to APOC2 overexpression in CRC

Our above findings have demonstrated the oncogenic role of APOC2 in CRC, still, how APOC2 could be aberrantly expressed in CRC remains unknown. Next, to identify the cause of APOC2 overexpression in CRC, we first performed DNA methylation analysis as well as copy number analysis of APOC2 promoter using the human methylation 450K BeadChip data from TCGA database. As it shown in Figure 8A and 8B, the mRNA expression of APOC2 is negatively correlated with APOC2 DNA methylation. Furthermore, analysis of APOC2 promoter DNA methylation reveals that the DNA methylation level of normal tissue is significantly higher than that in CRC tissues (Fig. 8C), indicating that DNA hypo-methylation of APOC2 promoter is one of the reasons for APOC2 overexpression in CRC.

**Fig. 8.**
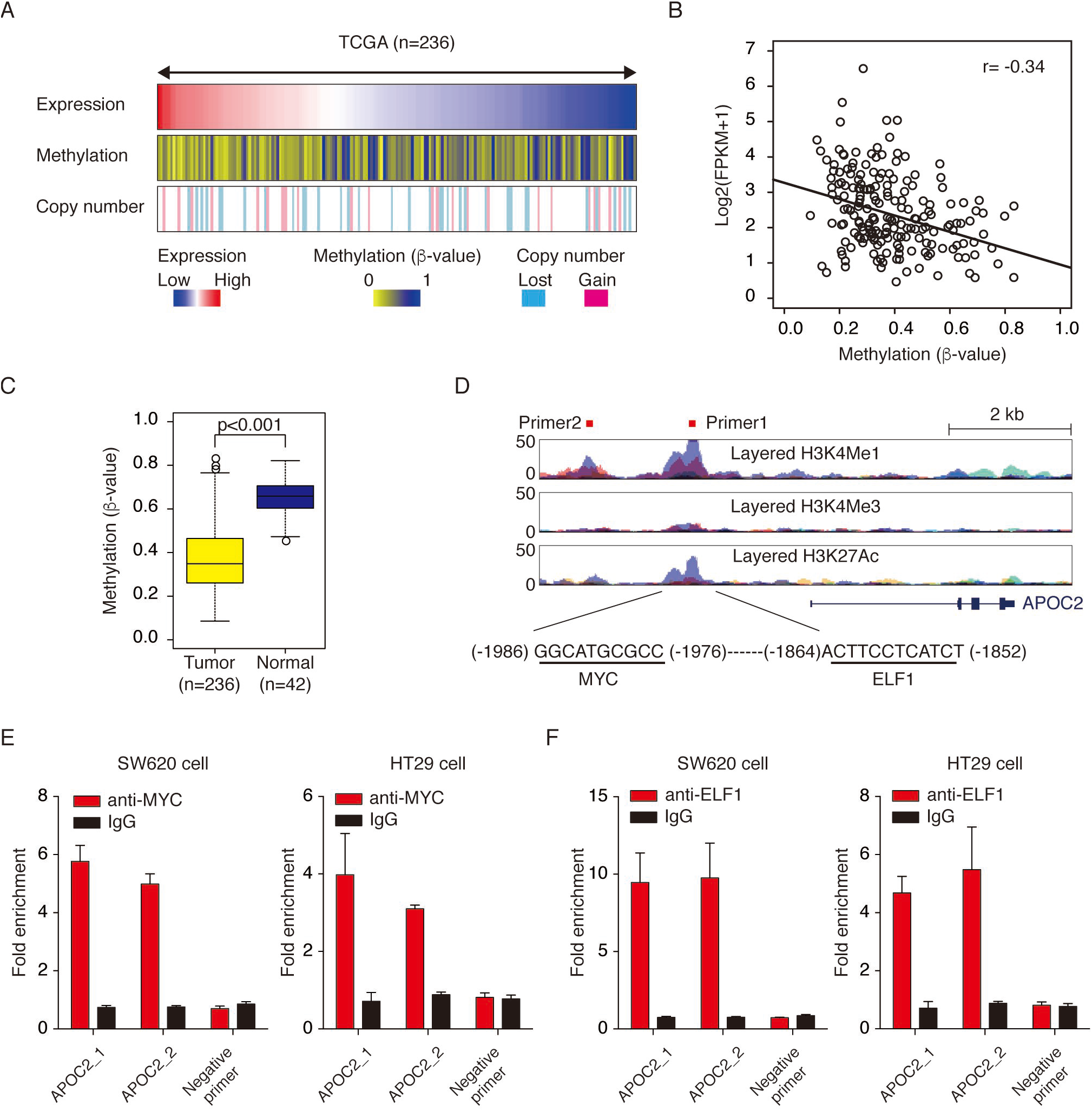
DNA hypo-methylation and the transcriptional activation by MYC and ELF1 contribute to APOC2 overexpression in CRC. (**A**) Evaluation of the association of copy number variation, APOC2 promoter methylation and APOC2 mRNA expression. The mRNA expression data were obtained from RNA-seq data of TCGA database. The DNA methylation data were obtained from human 450k methylation BeadChip of TCGA datasets. (**B**) DNA methylation of APOC2 negatively correlated with APOC2 mRNA expression. (**C**) The DNA methylation level of CRC tumor samples was significantly lower than normal tissues. The DNA methylation of 236 CRC samples and 42 normal tissue samples were obtained from TCGA database. (**D**) A representative image of regulatory element and motif in the promoter region of APOC2. (**E** and **F**) MYC and ELF1 bound at the promoter region of APOC2. ChIP-qPCR was performed in SW620 and HT29 cells.

Next, to identify the potential transcription factors that contribute to APOC2 overexpression, we integrated the transcription factor motifs identified by motif scan and the expression of its associated transcriptional factor, and found two potential transcription factor MYC and ELF1 might involve in APOC2 regulation (Fig. 8D). We thus performed the chromatin immunoprecipitation (ChIP) experiments using antibodies against MYC and ELF1, and validated the binding of these two transcription factors on the APOC2 promoter. As it shown in Figure 8E and 8F, both MYC and ELF1 bound at the promoter region of APOC2 in HT29 and SW620 cells. Together, our results indicating that in addition to DNA hypo-methylation, transcriptional activation by MYC and ELF1 also contribute to APOC2 overexpression.

## Discussion

Dysfunction of cellular lipid metabolism is one of the key biological processes during the progression of colorectal cancer. Here, we report that APOC2, an apolipoprotein, drives CRC proliferation and metastasis via lipid dysregulation in CRC cells. We demonstrated that APOC2 is overexpressed in CRC cells and its expression level was correlated with CRC metastasis. We found that APOC2 directly interacted with FASN and resulted in decreased anti-cancer metabolites such as EPA and DHA. The APOC2 overexpression also leads to upregulation of cellular levels of cellular α-KG, a cofactor required for activity of PHF8, a histone methylase which regulates transcription of genes involved in multiple pathways associated with tumorigenesis and metastasis.

Currently, the prognostic methods applied in clinical evaluation of CRC are the TNM staging system and measurement of levels of carcinoembryonic antigen (CEA)[29–31]. These methods work well but still have limitations, especially for patients diagnosed with stage II or III CRC. Thus, additional prognostic markers would be of great value. Many factors associated with prognosis have been intensively discussed, including lipoproteins as well as apolipoproteins. High levels of apolipoprotein B have been demonstrated to be a prognostic factor for poor outcomes in CRC. Low levels of apolipoprotein A1 in patients with CRC are associated with poor survival. Apolipoprotein C-I and apolipoprotein A2 have been considered as potential prognostic biomarkers for malignant mesothelioma and renal cancer, respectively. These studies indicate that apolipoproteins are essential for cancer progression and might serve as potential biomarkers for cancer prognosis. In our study, we demonstrate a positive correlation between APOC2 levels and poor prognosis of CRC. Our findings suggest that APOC2 might be applied as a biomarker for prognosis in CRC, although further evaluation in large clinical cohort studies is required.

Cancer cell metastasis is the major cause of therapy failure in CRC[32, 33]. Identification of the genes involved in CRC metastasis may help us to develop novel therapeutics strategies. In our study, we demonstrated that high levels of APOC2 were observed in the primary cancer tissue of patients with metastasis. In addition, a higher level of APOC2 expression was observed in metastasized tissues, indicating that APOC2 is an important factor in CRC metastasis. Moreover, both our *in vitro* invasion model and *in vivo* murine CRC model demonstrated that APOC2 directly contributes to CRC metastasis, and that decreasing APOC2 expression significantly inhibits CRC progression and metastasis. Together, our results demonstrate a role for APOC2 in CRC metastasis, and suggest that targeting APOC2 might be a potential therapeutic approach for CRC patients with metastasis.

Although the lipid transport role of apolipoproteins is well known, several studies also suggest that apolipoproteins play an additional role in regulating several biological processes which influence tumorigenesis and cancer progression. Apolipoprotein D binds to a cytokine receptor that mediates MAP kinase signaling thereby leading to cancer development[34]. Moreover, apolipoprotein B represses angiogenesis by modulating expression of VEGF receptor 1[17]. Cancer cell-secreted APOE functions as an angiogenesis and metastasis-suppressive factor through its association with distinct APOE receptors on melanoma and endothelial cells[16, 35]. Of note, these novel functions of apolipoproteins in cancer cells are exerted directly by the apolipoproteins themselves rather than through lipid moieties within the lipoprotein particles, further indicating that the role of apolipoproteins in tumorigenesis is far more complicated than expected. Thus, in addition to their functions in serum lipid transport, apolipoproteins may play versatile roles in tumorigenesis. In our study, we observed that overexpression of APOC2 accelerates cell proliferation and tumor cell invasion in CRC cell lines *in vitro* and *in vivo*. Inhibition of APOC2 expression inhibits cell growth as well as cell invasion in CRC cells and prolongs the survival of mice with CRC. Together, both *in vitro* and *in vivo* studies unveil an important oncogenic role for APOC2 in CRC.

Our functional studies indicate that APOC2-driven CRC progression and metastasis may be mediated by direct association of APOC2 with FASN, which may in turn result in decreased levels of anti-cancer metabolites such as EPA and DHA. FASN is the key protein that links cellular lipid metabolism to energy supply as well as to production of membrane lipids[36]. FASN has been implicated in the development of multiple cancers and is one of the most well-studied fatty acid metabolic enzymes in cancer[37, 38]. Aberrant expression of FASN correlates with poor prognosis of multiple cancers due to its role in *de novo* fatty acid synthesis[39]. FASN inhibition in APOC2 overexpressing cell lines suppresses tumorigenesis and metastasis, which indicates that APOC2-FASN-regulated lipid metabolism may contribute to CRC progression and metastasis. In addition, our studies indicated that APOC2 accelerates tumor progression by reducing docosahexaenoic acid (DHA) and eicosapentaenic acid (EPA) levels. Treatment with DHA and EPA inhibits growth of CRC cells. These results support the notion that dietary omega-3 fatty acids could benefit patients with colorectal cancer. DHA and EPA are two essential omega-3 polyunsaturated fatty acids which have been shown to exert anticancer effects in several cancer types, both *in vitro* and *in vivo*[40, 41]. EPA has also been shown to induce apoptosis in many tumor cells, including prostate cancer, breast cancer and medulloblastoma[42]. Dietary supplementation with omega-3 fatty acids is likely to prevent cancer[43]. In addition, studies on many animal models have shown that high doses of EPA and DHA improve the efficacy of chemotherapy while reducing the side effects. Our studies show for the first time that degradation of omega-3 fatty acids is associated with CRC tumorigenesis *in vitro*. We propose that APOC2 interacts with FASN to degrade EPA and DHA in CRC, and that this contributes to tumorigenesis and metastasis. However, the mechanisms by which APOC2 and FASN act together to degrade EPA and DHA remain unknown and require further investigation.

Metabolites generated from the process of lipid metabolism are the major sources for post-translational modifications including histone modification[44]. For the first time, we documented that increased α-KG levels are associated with CRC tumorigenesis *in vitro*. Increased α-KG levels result in activation of PHF8 and upregulated transcription of target genes. PHF8 is a JmjC domain-containing demethylase and demethylates mono- and dimethylated histone H3 Lys-9 residue (H3K9Me1 and H3K9Me2), dimethylated H3 Lys-27 (H3K27Me2), and monomethylated histone H4 Lys-20 residue (H4K20Me1)[45, 46]. Since H3K9Me1, H3K9Me2, H3K27Me2 and H4K20Me1 are generally epigenetic repressive loci, PFH8-mediated demethylation of these loci results in upregulation of downstream genes such as ACACA, DAD1, MAPK1 and NFKB1. Indeed, we observed that APOC overexpression is associated with the upregulation of those genes. Based on the results of our study, we propose that APOC2 affects α-KG levels, which leads to activation of PHF8. However, the mechanism by which APOC2 activates PHF8 remains elusive and warrants further investigation.

## Materials and Methods

### Patients and Tissue Samples

32 paired fresh frozen samples of primary colorectal carcinoma (CRC) and adjacent normal colon tissue were collected from the Department of Surgery at the Shanghai General Hospital, School of Medicine, Shanghai Jiaotong University. 507 paraffin-embedded samples of primary colorectal carcinoma were collected from two independent cohorts of CRC patients. Cohorts 1 and 2 comprised 276 patients diagnosed with CRC from 2003 to 2007 and 231 patients diagnosed from 2008 to 2011 at the Shanghai General Hospital, respectively. All samples were obtained during operations. None of these patients received any anti-cancer therapy before tumor resection. This research was approved by the Ethics Committee of Shanghai General Hospital, and written informed consents were obtained from all patients before being enrolled in the study.

### Tissue Microarray and Immunohistochemistry

Tissue microarrays were constructed using a tissue-arraying instrument (Shanghai Outdo Biotech Co., LTD, China). 507 paraffin-embedded tissue blocks were used for tissue microarray sampling. Representative 0.6-mm-diameter cylinders of tumor regions were punched from each tissue block and redeposited into a new paraffin block at predefined positions.

The intensity and extent of staining were evaluated independently by two pathologists who were blinded to patient outcomes. The staining intensity score was rated as 0 (no staining), 1 (mild staining), 2 (moderate staining), or 3 (intense staining). The area score was calculated as 0 (0%), 1 (1%–25%), 2 (26%–50%), 3 (51%–75%), or 4 (76%–100%) according to the percentage of positively-stained cells. The final scores were calculated by multiplying the intensity scores and area scores. Patients with CRC were divided into two groups according to the staining score: 0–8, lower expression; 9–12, higher expression.

### Cell culture, Plasmids, and Lentivirus

The Caco2, SW620, HT29, and RKO human colorectal cancer cells were purchased from American Type Culture Collection (Rockville, MD). All cells were cultured in Dulbecco’s modified Eagle medium (DMEM) (Gibco, USA) with 10% fetal bovine serum (FBS) (Gibco, USA). The APOC2 overexpression and knockdown lentivirus, as well as the vector virus, were purchased from Shanghai Genechem Co., Ltd., Shanghai, China.

### Chromatin-immunoprecipitation (ChIP)

Chromatin immunoprecipitation (ChIP) was performed in APOC2 overexpressing RKO cells or control RKO cells according to the manufacturer’s protocol of the Active Motif’s ChIP-IT High Sensitivity Kit (ActiveMotif, Carlsbad, CA) with modification. In brief, 1×10^7^ cells were fixed with 1% formaldehyde for 10 minutes in culture media at room temperature. The fixation reaction was quenched by adding 1/20 volume of 2.5 mM glycine for 5 minutes at room temperature. Cells were lysed in the 1 ml lysis buffer (10mM Tris-HCl PH 7.5, 10 mM NaCl, 3 mM MgCl2, 0.5% NP-40) for three times. The pellet was resuspended in ChIP buffer and sonicated for 20 cycles, 30s on and 30s off. For each ChIP, 5μg antibody for PHF8 (ab36068, Abcam) was added and incubated overnight at 4°C. 20μl Protein G beads were washed and added into the reaction for 3 hours. The beads were washed 5 times with buffer AM1. The immune complexes were eluted with 100 μL buffer AM4 at 37°C for 5 minutes and then 5μL proteinase K was added and incubated at 55°C for 30 minutes and 2 hours. The ChIP-seq data have been deposited at the GEO database with the following links: (https://www.ncbi.nlm.nih.gov/geo/query/acc.cgi?acc=GSE114089). The following secure token has been created to allow review of record GSE114089 while it remains in private status: ktqdoasofjanhwx

### Immunoblotting and Immunoprecipitation

Cells were washed 3 times with PBS before cell lysis. Cells were lysed in lysis buffer (20mM Tris-HCL (PH 8.0), 2mM EDTA, 1% Triton-X100, 150mM NaCl, 100 nM PMSF and 1x Proteinase Inhibitor cocktail). Antibodies against APOC2, FASN, GAPDH were used for immunoblotting. For immunoprecipitation, 2μg antibodies against APOC2(ab76452, Abcam) FASN (ab22759, Abcam) were added to cell lysate and incubated overnight at 4°C. 20μl Protein G beads (10003D, Invitrogen) were added to each reaction and incubated for 3 hours at 4°C. After washing 4 times with lysis buffer, the immunocomplex was eluted with elution buffer (25 mM Tris-HCl (PH 7.5), 10 mM EDTA, 1% SDS) followed by immunoblotting.

### RNA extraction and RT-qPCR

Total RNA was prepared using TRIzol reagent (Invitrogen, USA) according to the manufacturer’s instructions. First strand cDNA was synthesized from 2μg of total RNA using SuperScript® III First-Strand Synthesis System (Invitrogen, USA). Real-time quantification PCR was performed using the SYBR Premix Ex Taq II (Tli RNase H Plus) (Takara, Japan) with the following primers. APOC2 forward: 5’ CCCAGAACCTGTACGAGAAGAC 3’ and reverse 5’ AATGCCTGTGTAAGTGCTCATG3’. β-actin forward 5′-TCCTTAATGTCACGCACGATTT-3’ and reverse 5’-GAGCGCGGCTACAGCTT-3’. Each reaction was performed in triplicate.

### Cell Proliferation Assay

The CCK8 assay was used to evaluate the cell growth of APOC2 overexpressing RKO and Caco2 cells, APOC2 knockdown SW620 and HT29 cells, and all corresponding control cells. In brief, 1×10^4^ cells/well were seeded in 96-well plates containing 0.1 mL of medium. After 24 hours, 48 hours, 72 hours and 96 hours of cell culture, CCK8 solution (CK04-3000T, DoJindo, Japan) was added to each well (including the control well), and the mixture was incubated at 37°C for 2 hours. The 450-nm absorbance (OD450) of each well was measured by a microplate reader (Bio-Rad, USA).

### Wound Healing Assay

Cells were seeded in 6-well plates and allowed to grow to 70-80% confluence. Sterile white pipette tips were used to create wounds. Floating cells were removed by rinse two times with phosphate-buffered saline (PBS). Cells were cultured in DMEM supplemented with 10% FBS for 24 hours at 37°C. Cell migration was evaluated by measuring the relative percentage of wound region covered by migrated cells. Each assay was repeated 3 times.

### Migration and Invasion Assay

The migration and invasion assays were performed in 24-well Boyden chambers (Corning, NY) with filters (8-μm pore size) pre-coated with fibronectin (Roche). APOC2 overexpressing cells, APOC2 knockdown cells, or respective control cells were placed into the upper chamber with 0.5 ml serum-free DMEM (1×10^5^ cells per chamber). DMEM containing 10% fetal bovine serum was added to the lower chamber. After 24 hours, cells in the lower chamber were fixed with methanol for 5 minutes at room temperature followed by crystal violet staining. Experiments were repeated at least 3 times.

### In vivo tumor xenograft study

Four-week-old male nude mice were used in this study. For tumor growth evaluation, APOC overexpressing or knockdown cells, as well as respective control cells, were resuspended in serum-free DMEM, and 1×10^6^ APOC2 overexpressing or knockdown cells (in 0.1mL) were injected into the left flank of mice while their respective control cells were injected into the right flank. Tumor volumes were evaluated every week after injection. Tumors were removed and fixed in 10% formalin. Tumor volume and weight were measured.

For the lung metastasis model, 1×10^6^ cells (in 0.2 ml) stably expressing luciferase were injected into the tail vein of nude mice. For portal circulation injections, cells (10^6^/ml, 200 μl) were injected into the spleen followed by removal of the spleen. Animals were excluded from studies if inoculated cells did not arrive in the liver. *In vivo* bioluminescence was generated using d-luciferin substrate (Perkin Elmer) and bioluminescence signal was normalized to the signal detected immediately following cell inoculation.

### Statistical analysis

The Kaplan-Meier survival analysis was performed using the GraphPad Prism version 7.0a software. Statistical significance of the difference between survival curves for APOC2 high-expressing and low-expressing patients was assessed using log-rank tests. The significance of APOC2 expression between different groups was assessed using the unpaired or paired students’ t-test (2-tailed).

## Abbreviations

APOC2: apolipoprotein C2
CRC: Colorectal cancer
TCGA: The Cancer Genome Atlas
LC/MS: liquid chromatography–mass spectrometry
ChIP: Chromatin immunoprecipitation
EPA: eicosapentaenoic acid:
DHA: docosahexaenoic acid
α-KG: alpha-ketoglutarate acid
GSEA: Gene Set Enrichment Analysis
HR: Hazard ratio
IHC: Immunohistochemistry

## Acknowledgements

This study was supported by funds from Natural Science Foundation of China (NSFC) (81220108021, 81602069 and 81702337), and Project of Shanghai Science and Technology Commission (14411950502).

## Authors’ contributions

Jian Chen, Huamei Tang and Zhihai Peng designed this study; Zhihai Peng, Jian Chen and Yupeng Wang obtained fundings; Jian Chen and Chao Xiao performed the clinical studies. Jian Chen, Guohe Song, Chao Xiao, Yupeng Wang, Xueni Liu, Jiayi Chen and Jing Kuai performed the experiments. Jian Chen, Chao Xiao and Yupeng Wang wrote the manuscript, Xuebin Qin, Huamei Tang, Weiping Guo and Zhihai Peng revised the manuscript. All authors approved the final version of the manuscript.

## Conflict of interest

The authors have declared that no competing interest exists.

## References

1 Bijlsma MF, Sadanandam A, Tan P, Vermeulen L. Molecular subtypes in cancers of the gastrointestinal tract. Nat Rev Gastroenterol Hepatol 2017; 14: 333–342.

2 Siegel R, Desantis C, Jemal A. Colorectal cancer statistics, 2014. CA Cancer J Clin 2014; 64: 104–117.

3 Wu Z, Wei D, Gao W, Xu Y, Hu Z, Ma Z et al. TPO-Induced Metabolic Reprogramming Drives Liver Metastasis of Colorectal Cancer CD110+ Tumor-Initiating Cells. Cell Stem Cell 2015; 17: 47–59.

4 Beloribi-Djefaflia S, Vasseur S, Guillaumond F. Lipid metabolic reprogramming in cancer cells. Oncogenesis 2016; 5: e189.

5 Carracedo A, Cantley LC, Pandolfi PP. Cancer metabolism: fatty acid oxidation in the limelight. Nat Rev Cancer 2013; 13: 227–232.

6 Mancini R, Noto A, Pisanu ME, De Vitis C, Maugeri-Sacca M, Ciliberto G. Metabolic features of cancer stem cells: the emerging role of lipid metabolism. Oncogene 2018; 37: 2367–2378.

7 Phipps AI, Limburg PJ, Baron JA, Burnett-Hartman AN, Weisenberger DJ, Laird PW et al. Association between molecular subtypes of colorectal cancer and patient survival. Gastroenterology 2015; 148: 77–87 e72.

8 Boroughs LK, DeBerardinis RJ. Metabolic pathways promoting cancer cell survival and growth. Nat Cell Biol 2015; 17: 351–359.

9 Rocha CM, Carrola J, Barros AS, Gil AM, Goodfellow BJ, Carreira IM et al. Metabolic signatures of lung cancer in biofluids: NMR-based metabonomics of blood plasma. J Proteome Res 2011; 10: 4314–4324.

10 Martin LJ, Melnichouk O, Huszti E, Connelly PW, Greenberg CV, Minkin S et al. Serum lipids, lipoproteins, and risk of breast cancer: a nested case-control study using multiple time points. J Natl Cancer Inst 2015; 107.

11 Ko HL, Wang YS, Fong WL, Chi MS, Chi KH, Kao SJ. Apolipoprotein C1 (APOC1) as a novel diagnostic and prognostic biomarker for lung cancer: A marker phase I trial. Thorac Cancer 2014; 5: 500–508.

12 Harima Y, Ikeda K, Utsunomiya K, Komemushi A, Kanno S, Shiga T et al. Apolipoprotein C-II is a potential serum biomarker as a prognostic factor of locally advanced cervical cancer after chemoradiation therapy. Int J Radiat Oncol Biol Phys 2013; 87: 1155–1161.

13 Chung L, Moore K, Phillips L, Boyle FM, Marsh DJ, Baxter RC. Novel serum protein biomarker panel revealed by mass spectrometry and its prognostic value in breast cancer. Breast Cancer Res 2014; 16: R63.

14 Vardy JL, Dhillon HM, Pond GR, Rourke SB, Bekele T, Renton C et al. Cognitive Function in Patients With Colorectal Cancer Who Do and Do Not Receive Chemotherapy: A Prospective, Longitudinal, Controlled Study. J Clin Oncol 2015; 33: 4085–4092.

15 Borgquist S, Butt T, Almgren P, Shiffman D, Stocks T, Orho-Melander M et al. Apolipoproteins, lipids and risk of cancer. Int J Cancer 2016; 138: 2648–2656.

16 Pencheva N, Tran H, Buss C, Huh D, Drobnjak M, Busam K et al. Convergent multi-miRNA targeting of ApoE drives LRP1/LRP8-dependent melanoma metastasis and angiogenesis. Cell 2012; 151: 1068–1082.

17 Avraham-Davidi I, Ely Y, Pham VN, Castranova D, Grunspan M, Malkinson G et al. ApoB-containing lipoproteins regulate angiogenesis by modulating expression of VEGF receptor 1. Nat Med 2012; 18: 967–973.

18 Takano S, Yoshitomi H, Togawa A, Sogawa K, Shida T, Kimura F et al. Apolipoprotein C-1 maintains cell survival by preventing from apoptosis in pancreatic cancer cells. Oncogene 2008; 27: 2810–2822.

19 Jong MC, Hofker MH, Havekes LM. Role of ApoCs in lipoprotein metabolism: functional differences between ApoC1, ApoC2, and ApoC3. Arterioscler Thromb Vasc Biol 1999; 19: 472–484.

20 Wolska A, Dunbar RL, Freeman LA, Ueda M, Amar MJ, Sviridov DO et al. Apolipoprotein C-II: New findings related to genetics, biochemistry, and role in triglyceride metabolism. Atherosclerosis 2017; 267: 49–60.

21 Kei AA, Filippatos TD, Tsimihodimos V, Elisaf MS. A review of the role of apolipoprotein C-II in lipoprotein metabolism and cardiovascular disease. Metabolism 2012; 61: 906–921.

22 Xue A, Chang JW, Chung L, Samra J, Hugh T, Gill A et al. Serum apolipoprotein C-II is prognostic for survival after pancreatic resection for adenocarcinoma. Br J Cancer 2012; 107: 1883–1891.

23 Brown I, Cascio MG, Rotondo D, Pertwee RG, Heys SD, Wahle KW. Cannabinoids and omega-3/6 endocannabinoids as cell death and anticancer modulators. Prog Lipid Res 2013; 52: 80–109.

24 Cockbain AJ, Toogood GJ, Hull MA. Omega-3 polyunsaturated fatty acids for the treatment and prevention of colorectal cancer. Gut 2012; 61: 135–149.

25 Cockbain AJ, Volpato M, Race AD, Munarini A, Fazio C, Belluzzi A et al. Anticolorectal cancer activity of the omega-3 polyunsaturated fatty acid eicosapentaenoic acid. Gut 2014; 63: 1760–1768.

26 Loenarz C, Schofield CJ. Expanding chemical biology of 2-oxoglutarate oxygenases. Nat Chem Biol 2008; 4: 152–156.

27 Ren R, Ocampo A, Liu GH, Izpisua Belmonte JC. Regulation of Stem Cell Aging by Metabolism and Epigenetics. Cell Metab 2017; 26: 460–474.

28 Chowdhury R, Sekirnik R, Brissett NC, Krojer T, Ho CH, Ng SS et al. Ribosomal oxygenases are structurally conserved from prokaryotes to humans. Nature 2014; 510: 422–426.

29 Tiernan JP, Perry SL, Verghese ET, West NP, Yeluri S, Jayne DG et al. Carcinoembryonic antigen is the preferred biomarker for in vivo colorectal cancer targeting. Br J Cancer 2013; 108: 662–667.

30 Hanash SM, Baik CS, Kallioniemi O. Emerging molecular biomarkers--blood-based strategies to detect and monitor cancer. Nat Rev Clin Oncol 2011; 8: 142–150.

31 Stern E, Vacic A, Rajan NK, Criscione JM, Park J, Ilic BR et al. Label-free biomarker detection from whole blood. Nat Nanotechnol 2010; 5: 138–142.

32 Chambers AF, Groom AC, MacDonald IC. Dissemination and growth of cancer cells in metastatic sites. Nat Rev Cancer 2002; 2: 563–572.

33 Wirtz D, Konstantopoulos K, Searson PC. The physics of cancer: the role of physical interactions and mechanical forces in metastasis. Nat Rev Cancer 2011; 11: 512–522.

34 Soiland H, Soreide K, Janssen EA, Korner H, Baak JP, Soreide JA. Emerging concepts of apolipoprotein D with possible implications for breast cancer. Cell Oncol 2007; 29: 195–209.

35 Lee JH, Giannikopoulos P, Duncan SA, Wang J, Johansen CT, Brown JD et al. The transcription factor cyclic AMP-responsive element-binding protein H regulates triglyceride metabolism. Nat Med 2011; 17: 812–815.

36 Wei X, Song H, Yin L, Rizzo MG, Sidhu R, Covey DF et al. Fatty acid synthesis configures the plasma membrane for inflammation in diabetes. Nature 2016; 539: 294–298.

37 Rohrig F, Schulze A. The multifaceted roles of fatty acid synthesis in cancer. Nat Rev Cancer 2016; 16: 732–749.

38 Yang CS, Matsuura K, Huang NJ, Robeson AC, Huang B, Zhang L et al. Fatty acid synthase inhibition engages a novel caspase-2 regulatory mechanism to induce ovarian cancer cell death. Oncogene 2015; 34: 3264–3272.

39 Li CF, Fang FM, Chen YY, Liu TT, Chan TC, Yu SC et al. Overexpressed Fatty Acid Synthase in Gastrointestinal Stromal Tumors: Targeting a Progression-Associated Metabolic Driver Enhances the Antitumor Effect of Imatinib. Clin Cancer Res 2017; 23: 4908–4918.

40 Fluckiger A, Dumont A, Derangere V, Rebe C, de Rosny C, Causse S et al. Inhibition of colon cancer growth by docosahexaenoic acid involves autocrine production of TNFalpha. Oncogene 2016; 35: 4611–4622.

41 Chung H, Lee YS, Mayoral R, Oh DY, Siu JT, Webster NJ et al. Omega-3 fatty acids reduce obesity-induced tumor progression independent of GPR120 in a mouse model of postmenopausal breast cancer. Oncogene 2015; 34: 3504–3513.

42 Torres-Adorno AM, Vitrac H, Qi Y, Tan L, Levental KR, Fan YY et al. Eicosapentaenoic acid in combination with EPHA2 inhibition shows efficacy in preclinical models of triple-negative breast cancer by disrupting cellular cholesterol efflux. Oncogene 2018.

43 Vaughan VC, Hassing MR, Lewandowski PA. Marine polyunsaturated fatty acids and cancer therapy. Br J Cancer 2013; 108: 486–492.

44 Wong CC, Qian Y, Yu J. Interplay between epigenetics and metabolism in oncogenesis: mechanisms and therapeutic approaches. Oncogene 2017; 36: 3359–3374.

45 Yu L, Wang Y, Huang S, Wang J, Deng Z, Zhang Q et al. Structural insights into a novel histone demethylase PHF8. Cell Res 2010; 20: 166–173.

46 Horton JR, Upadhyay AK, Qi HH, Zhang X, Shi Y, Cheng X. Enzymatic and structural insights for substrate specificity of a family of jumonji histone lysine demethylases. Nat Struct Mol Biol 2010; 17: 38–43.

